# Bacteria.guru: comparative transcriptomics and co-expression database for bacterial pathogens

**DOI:** 10.1101/2021.08.02.454836

**Authors:** Peng Ken Lim, Emilia E Davey, Sean Wee, Wei Song Seetoh, Jong Ching Goh, Xinghai Zheng, Sean Kia Ann Phang, Eugene Sheng Kai Seah, Janice Wan Zhen Ng, Xavier Jia Hui Wee, Aloysius Jun Hui Quek, Jordan JingHeng Lim, Edbert Edric Rodrigues, Heesoo Lee, Chin Yong Lim, Wei Zhi Tan, Yuet Ruh Dan, Bronson Lee, Samuel En Le Chee, Zachary Ze En Lim, Jia Sheng Guan, Ivan Jia Le Tan, Trinidad Jeremiah Arong, Marek Mutwil

**Affiliations:** School of Biological Sciences, Nanyang Technological University, 60 Nanyang Drive, Singapore, 637551, Singapore

## Abstract

**Summary:** The bacterial kingdom comprises unicellular prokaryotes able to establish symbioses from mutualism to parasitism. To combat bacterial pathogenicity, we need an enhanced understanding of gene function and regulation, which will mediate the development of novel antimicrobials. Gene expression can predict gene function, but there lacks a database enabling expansive inter- and intraspecific exploration of gene expression profiles and co-expression networks for bacteria. To address this, we integrated the genomic and transcriptomic data of the 17 most notorious and studied bacterial pathogens, creating bacteria.guru, an interactive database that can identify, visualize, and compare gene expression profiles, co-expression networks, functionally enriched clusters, and gene families across species. Through illustrating antibiotic resistance mechanisms in *P. aeruginosa*, we demonstrate that bacteria.guru could potentially aid the discovery of multi-faceted antibiotic targets. Hence, we believe bacteria.guru will facilitate future bacterial research.

**Availability:** The database and co-expression networks are freely available from https://bacteria.guru/. The sample annotations are found in the supplemental data.

## Introduction

Bacteria are ubiquitous unicellular organisms that constitute a diverse kingdom. Despite their fundamental role in the biosphere, the synergistic relationship between the rapid evolution and acute virulence of the pathogenic variety poses the urgency for novel antimicrobials (Davies and Davies, 2010), especially considering the negligent use of antibiotics. The combination of these factors precipitated an epidemic of antibiotic resistance; for example, following the introduction of □-lactams, which inhibit cell wall peptidoglycan synthesis, many bacterial pathogens increased the production of □-lactamases, rendering these medications ineffective. As bacteria continue to evolve these novel resistance mechanisms, our inability to combat them becomes increasingly exposed.

Several methodologies can be used to study genes essential for virulence and antibiotic resistance, such as sequence similarity, gene expression, and co-expression network analyses (Van dam et al., 2018; Jiang et al., 2016). Since predicting gene function from gene sequence alone can result in only a partial or incorrect prediction (Gerlt and Babbitt, 2000; Rhee and Mutwil, 2014), other methods to deduce gene function have emerged. Co-expression analysis is based on the observation that genes which exhibit similar expression profiles across different growth conditions and genotypes are often functionally related (Mutwil et al., 2011), and the utility of co-expression networks has exponentiated our understanding of gene function in multiple kingdoms of life. These networks comprise genes transcriptionally coordinated across various environmental conditions and growth stages, wherein nodes correspond to genes, and edges represent their significant co-expression relationships (Ruan et al., 2010). Co-expression networks are now used to impart insight into the evolution and interspecific conservation of gene modules (Ruprecht et al., 2017), gene function, gene regulation, and subcellular localization of gene products (Mutwil et al., 2011).

Yet, there lacks a platform enabling expansive exploration of inter-and intraspecific gene expression for bacteria. To address this, we present bacteria.guru, a database comprising RNA-sequencing data for 17 notorious bacterial pathogens. In addition to identifying and analyzing co-expression networks and conserved gene modules, bacteria.guru enables the within-and across species analysis of gene expression profiles, gene families, ontology terms, and possible gene-cluster redundancy.

## Implementation

Bacteria.guru is based on the CoNekT framework (Proost and Mutwil, 2018). The database supplies gene pages, each containing functional annotations, cDNA and protein sequences, the assigned gene family and corresponding phylogenetic tree, expression profiles, the co-expression neighborhood and cluster, significantly similar neighborhoods, and Gene Ontology information (https://bacteria.guru/features). For a description of the exhaustive construction process of bacteria.guru, refer to the Supplementary Methods.

To demonstrate how our database can be used to generate novel insights, we use *Pseudomonas aeruginosa* (*P. aeruginosa*) as a case study. Among the most common ICU-acquired infections in critically ill patients (Pachori et al., 2019), the pathogen is not only multidrug-resistant but was also recently discovered to have adaptive resistance capabilities (Taylor et al., 2014). To better comprehend antibiotic resistance in *P. aeruginosa*, ‘antibiotic metabolic process’ (GO:0016999) was queried using the ‘Find enriched clusters’ tool (https://bacteria.sbs.ntu.edu.sg/search/enriched/clusters), where a cluster containing 238 genes, cluster 12 (https://bacteria.sbs.ntu.edu.sg/cluster/view/467), was identified to be significantly enriched (p-value < 0.05). Analysis of Cluster 12 revealed gene members involved in several key processes, which include phenazine synthesis, and quorum sensing (QS), alongside antibiotic metabolism (Fig. 1B), which implicates these processes in *P. aeruginosa* resistance mechanisms. Production of phenazine has been proposed to alter antibiotic susceptibility (Schiessl et al., 2019) and affect swarming motility (Ramos et al., 2010) at early stages of biofilm formation, whereas QS serves as an important regulatory mechanism for coordinating biofilm formation (Subramani et al., 2019). By exploring these interrelations, we could determine whether adaptive antibiotic resistance is achieved through the mechanism of biofilm aggregation. The presence of several proteins related to flagellum-hooking, fimbrial assembly, and cell-adhesion in the cluster, which are essential in biofilm formation (Haiko and Westerlund-Wikström, 2013; Ruer et al., 2007), supports this premise; in fact, Miller and colleagues observed biofilm-induction by aminoglycoside antibiotics in *P. aeruginosa* (Hoffman et al., 2005), demonstrating how our database can be used to unravel potential antibiotic-induced epigenetic changes.

**Figure 1.**
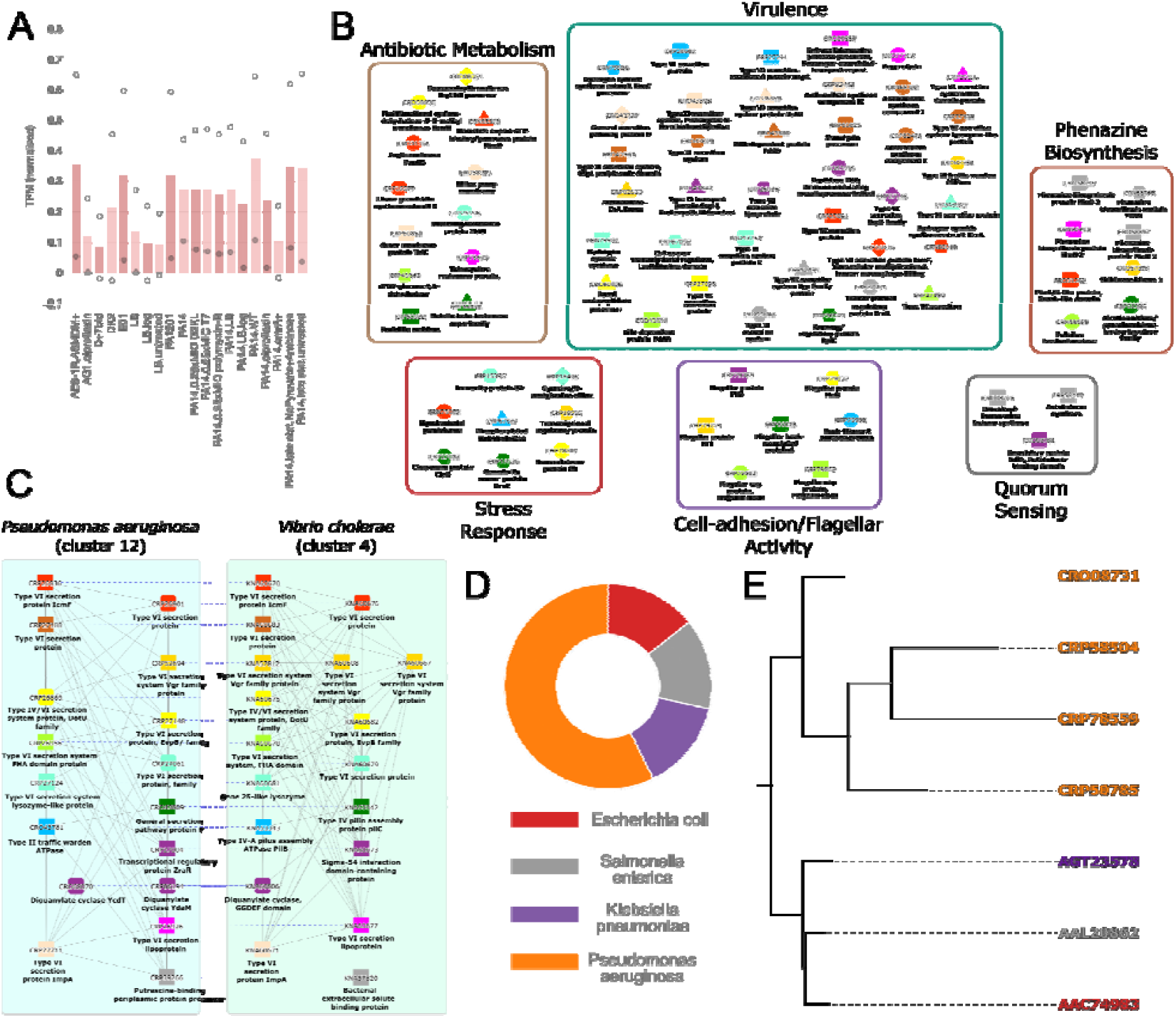
Antibiotic metabolic process analysis in *Pseudomonas aeruginosa*. A) Average expression profile snippet of cluster 12 for *P. aeruginosa*. The x-axis and y-axis represent different samples (capturing genotype and growth condition) and gene expression levels (as transcript per million), respectively. The bars indicate the average expression, while the points represent the maximum/minimum expression in a given experiment. B) Co-expression cluster 12 from P. aeruginosa enriched for GO:0016999 (Antibiotic Metabolic Process). Nodes, colored shapes, and gray edges represent genes, orthogroups, and co-expressed genes, respectively. For concision, only co-expressed genes involved in the discussed processes are shown. C) Conserved clusters involved in virulence in *Pseudomonas aeruginosa* (cluster 12, left) and *Vibrio cholerae* (cluster 4, right). Dashed edges connect genes belonging to the same orthogroup. D) The donut chart indicates the proportion of genes belonging to *Pseudomonas aeruginosa* (4 genes), *Escherichia coli* (1 gene), *Klebsiella pneumoniae* (1 gene), and *Salmonella enterica* (1 gene) in the OG_02_0001636 orthogroup. E) Gene tree of the OG_02_0001636 orthogroup (*P. aeruginosa* paralogs comprise the upper clade). The color coding between the donut charts and gene trees is conserved.

Using the ‘similar clusters’ tool, one can identify similar clusters in other bacterial species for further analysis. Here we identified and compared a similar cluster in *Vibrio cholerae* to cluster 12, with a Jaccard index of 0.062 (Fig. 1C). To illustrate the database’s ability to provide ortholog information on genes, a gene encoding for SdiA, an autoinducer receptor, from cluster 12 (ID: CRO08731) was selected. Clicking on the gene’s family (<u>OG_02_0001636</u>) located on the gene page (Fig. 1D), followed by the corresponding phylogenetic tree link, disclosed orthologs found in three other bacterial species (Fig. 1E). Furthermore, the presence of four *sdiA* paralogs denotes a critical role of QS in *P. aeruginosa* antibiotic resistance. With the interplay of QS in formation and regulation of resistance mechanisms necessitating further study (Zhao et al., 2020), additional scrutiny of this biological process overlap could illuminate the nexus of microbial resistance.

## Conclusion

The need to better comprehend antibiotic resistance motivated our construction of bacteria.guru. The database offers inter-and intraspecific analyses of co-expressed gene networks and neighbourhoods, expression profiles, gene families, phylograms, and ontology terms in 17 pathogens. We demonstrated how bacteria.guru can be used to study *P. aeruginosa* antibiotic metabolism, but we envision that its function can be extended to explore any biological pathway in these bacteria.

## SUPPLEMENTARY INFORMATION

**Supplementary Methods 1**. Tools and analyses used to construct bacteria.guru

### Supplementary tables

Table S1. Genome versions used to build the database

Table S2. Annotations of the RNA-seq samples used in the study

