## Supplemental Methods 1 for "Bacteria.guru: comparative transcriptomics and co-expression database for bacterial pathogens"

Coding sequences (CDSs) (Table S1) and gene expression data of the 17 species were obtained from a recent crowd-sourced analysis of bacterial RNA sequencing data (Hew et al., 2020), where gene expression was quality-controlled (Table S2) and quantified via Kallisto pseudoalignment (Bray et al., 2016). To identify orthogroups, CDSs were fed into OrthoFinder v2.312 (Emms and Kelly, 2015) using Diamond (Buchfink et al., 2015) with default settings, and the FastTree algorithm (Price et al., 2009) was used to construct gene trees. To identify Pfam domains and Gene ontology (GO) terms, we used the onboard conversion function of CoNekt (Proost and Mutwil, 2018) to generate PEP files from CDSs, and subjected them to Interproscan-5.51-85.0-5.44-79 (Jones et al., 2014) analysis. For each species, Pearson Correlation Coefficients (PCC) of gene pairs were calculated based on expression across experiments. Highest Reciprocal Rank (HRR) (Mutwil et al., 2010) was used to construct co-expression networks, and co-expression clusters were generated via Heuristic Cluster Chiseling Algorithm (HCCA) (Mutwil et al., 2010) with a cluster size of 100 genes. Experiment annotations (Table S20), including the specific bacterial strains and growth perturbations, were derived from SRA runtables downloaded from the NCBI (National Center for Biotechnology Information) database, and were used to construct the expression profiles. Makeblastdb v2.9.0+ (Camacho et al., 2009) was used to construct the protein and nucleotide blast databases for the BLAST search functionality featured on bacteria.guru.
